# Modelling the dynamics of transposable elements in genomes under asexual reproduction using agent-based model

**DOI:** 10.1101/2025.09.28.679026

**Authors:** Martin Rosalie, Moaine El Baidouri, Sébastien Gourbière

**Affiliations:** UMR5096 Laboratoire Génome et Développement des Plantes, Université de Perpignan Via Domitia, CNRS, Perpignan, France; Centre for the Study of Evolution, School of Life Sciences, University of Sussex, Brighton BN1 9QG, United Kingdom

**Keywords:** Transposition regulation, Model, Computer simulation, Genome evolution, Gillespie algorithm, Asexual reproduction

## Abstract

Transposable elements (TEs) are abundantly present in eukaryotic genomes and can be likened to parasites colonizing a genome due to their properties. From this perspective, a population-based approach has been developed to model interactions between TEs and a genome population. The distribution of TEs within a population of genomes is studied over the long term to understand the mechanisms allowing TEs to persist despite their deleterious effects on genomes. Under this single restrictive assumption, the results show that the population of TEs can persist for a very long time within the genome population, when the genome population is highly diverse in terms of the distribution of TEs quantities. When there is no mechanism for silencing TEs, the proposed model of asexual reproduction either purges TEs or leads to co-extinction of genomes and TEs. On the other hand, with a high proportion of silenced TEs, the population of TEs can be maintained for a long time in the population of genomes.

## 1. Introduction

Transposable elements (TEs) are mobile genetic elements able to replicate and transpose within the genome, leading to an increase in their copy number. TEs constitute a substantial proportion of eukaryotic genomes causing genome size variations (Tenaillon et al., 2010; Deragon et al., 2008; Wicker et al., 2018). Given their selfish nature and their abilities to disrupt coding genes leading to deleterious effects, TEs are seen as parasites that exploit host resources to sustain their own population (Burt and Trivers, 2008). Far from being “junk DNA”, TEs have shaped the evolution of gene regulation and genome architecture (Feschotte, 2008). They can act as promoters (Lai et al., 2009; Flemr et al., 2013; Butelli et al., 2012) and cis-regulatory elements (Xie et al., 2023; Lu et al., 2019) that influence the expression of neighboring genes. They can also affect alternative splicing (Varagona et al., 1992; Berthelier et al., 2023) and cause genome rearrangements such as deletions and duplications (Ma et al., 2023), among many other functional and structural impacts. As a result, TEs activation has been associated with species adaptation and evolution (Catoni, 2024) and is a source of important phenotypic variations (Tao et al., 2025).

Various population genetics models simulate and reproduce the behaviour of these TEs within genomes (Le Rouzic and Capy, 2005; Le Rouzic et al., 2007). Host genomes have evolved efficient mechanisms to silence TE activity through epigenetic regulation. These responses reduce the transpositional activity of TEs, thereby mitigating the risks of their invasion into the genome. These include DNA methylation (Deniz et al., 2019), histone modifications (Pachamuthu and Borges, 2023; Lık et al., 2025), and small RNA-based targeting, such as piRNAs in mammals (Ernst et al., 2017) and siRNAs in plants (Ito, 2011; Roessler et al., 2018). The former mechanisms involving clusters trigger a defensive response after an accumulation of TEs in these genomic regions and theoretical models exists (Kofler, 2019; Tomar et al., 2023). In addition to population genetics models, population-based models allow for alternative theoretical predictions for asexual reproduction and assuming deleterious effect of TEs (Flores-Ferrer et al., 2021). Building on these theoretical predictions, we conducted individual-based numerical simulations to study the structure of the distribution of TEs and its influence on their dynamics.

## 2. Models and methods

### 2.1 Biological system

Copy-and-paste TEs, also known as retrotransposons, propagate via a RNA intermediate that undegoes reverse transcription. Indeed, these elements are first transcribed into RNA, which is then reverse-transcribed into complementary DNA (cDNA) by reverse transcriptase. The resulting cDNA is subsequently integrated into a new genomic location by an integrase enzyme. This process leads to the accumulation of multiple copies of the element throughout the genome. Retrotransposons include LINEs, SINEs, and LTR elements, and constitute a major source of genome expansion and variability. However, their mobility also poses a threat to genome stability. Insertional mutagenesis, disruption of coding or regulatory regions, and chromosomal rearrangements are potential deleterious outcomes of unchecked TE activity (Hancks and Kazazian, 2012). As a result, organisms have evolved sophisticated epigenetic silencing mechanisms, such as the piRNA pathway, to control TE mobilization, particularly in germ cells (Siomi et al., 2011). Thus, TEs represent both a source of evolutionary innovation and a genomic burden that is tightly regulated. In parallel, theoretical models developed by Le Rouzic and Capy have explored the population-level dynamics of TEs, highlighting the balance between transposition, selection, and drift in shaping TEs content across genomes (Le Rouzic and Capy, 2005, 2006; Le Rouzic et al., 2007; Tomar et al., 2023). These models emphasize how TE-host interactions evolve under both genetic and ecological constraints. In addition to modeling, genomic comparisons can be performed, such as those by (Banuelos and Sindi, 2018), who modeled the density of complete and partial copies within host genomes and compared the resulting TE length distributions with annotated genomic collections.

Given their properties, TEs are modeled as parasites that exploit the opportunities offered by their hosts, the genomes (Hickey, 1982). Within this framework, theoretical models describing TE dynamics under the assumption of asexual reproduction (Charlesworth and Charlesworth, 1983; Dolgin and Charlesworth, 2006) all indicate that only two long-term outcomes are possible: either an explosion in the number of TEs leading to host extinction, or the elimination of TEs. However, recent examples exist of species reproducing asexually that still harbor recently active TEs (Bast et al., 2015). In rotifers, it has been shown (Nowell et al., 2021) that the expansion of RNAi gene-silencing pathways has enhanced their defense against the deleterious effects of TEs.

### 2.2 Eco–genomic model

A mathematical model describing the interactions between population of genomes and TEs for asexual reproduction has been proposed (Flores-Ferrer et al., 2021). This model improves a previous model (May and Anderson, 1978) that consider a finite number of asexual hosts carrying an infinite number of parasites, inducing for our purpose: genomes carrying an infinite number of TEs. The infinite set of differential equations describing the (Flores-Ferrer et al., 2021) model are defined in (1) where *G*_*i,j*_(*t*) is the number of genomes with *i* active TEs and *j* silent TEs.

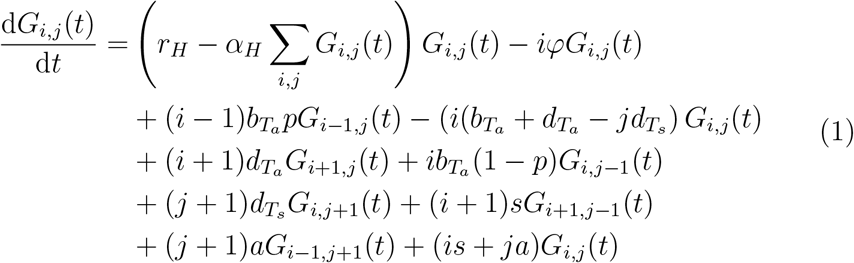

In this model, *r*_*H*_ = *b*_*H*_ − *d*_*H*_ refers to the reproduction rate of the genome (host in host-parasite formalism), *α*_*H*_ refers to the competition rate between genomes, *φ* is the deleterious effect of an active TE on the genome, *a* refers to the activation of a TE and reciprocally *s* refers to the silencing of a TE. Finally, 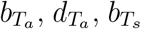 and 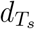 refer to birth and death rates of active and silent TEs. One of the main results of the latter article is that, with assumptions on the distribution of TEs over the genomes, there are some cases where the invasion of TEs is successful and maintainable without selective purge of TEs nor co-extinction of hosts and TEs populations.

The predictions of this mathematical model provide an overview scenario of TEs evolution in genomes and propose some conditions required for a successful invasion of TEs. We are addressing the specific case where the host growth is regulated (*α*_*H*_≠ 0). For this specific case, Fig. 1 details that potentially three scenarios can occur:

**Figure 1:**
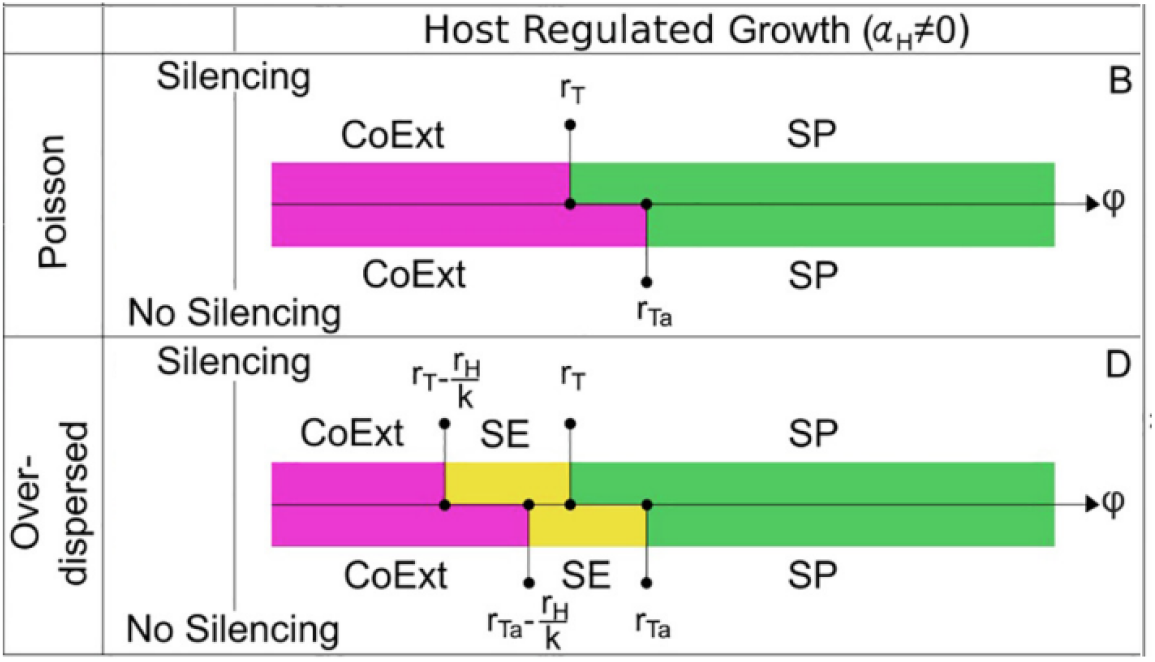
Reproduction of a portion of Fig. 4 of the article (Flores-Ferrer et al., 2021) with the following caption: “Conditions for the stabilization of TE-Host dynamics in the presence of TE copy number variations. The conditions where the dynamics presented in Fig. 3 occur are given with respect to *φ* under the assumptions considered for the host demography, the distribution of TEs, and with or without silencing. CoExt = Co-Extinction, DD = Demographic Dilution, SP = Selective Purge, SG = Stable Growth, SE = Stable coexistence Equilibrium”. The missing part of the original plot is related to the case where *α* = 0 leading to a host exponential growth that is not addressed here.

**CoExt: Co-Ext**inction occurs whenever there is an extinction of several species that is an aftermath result of the extinction of another species that their survival relied on. In that case of invasion, it means that the number of TEs increase and TEs fully invade the genomes until they become so numerous that the host cannot support their cumulative deleterious effects causing the decline of genome population followed by the TEs populations which no longer has hosts.

**SE: S**table coexistence **E**quilibrium represents a state where it is possible for several species to be able to coexist with each other over time without having one species completely driving the others to extinction. This refers to a successful invasion enabling TEs persistence.

**SP: S**elective **P**urge refers to a scenario where deleterious effects are so lethal that TEs cannot spread because of genomes. TEs are gradually eliminated, leading to their disappearance within the host population.

We reproduce here (Fig. 1) a portion of the figure of the initial article (Flores- Ferrer et al., 2021) summarising these results obtained via a mathematical analysis of differential equation systems presupposing on the distribution of TEs within the genome population. As the model is an analogy between parasites and TEs, the *φ* parameter represents the deleterious effect of a TE in genomes. The increase of *φ* lead to selective purge (SP) from co-extinction (Co-Ext) inducing that, theoreticaly, there is a range of parameters for *φ* leading to stable equilibrium (SE) where TEs survive to genomes population. These analytical results are obtained if, and only if, the distribution of the TEs follows an over-dispersed distribution (also known as a negative binomial distribution). This particular scenario is not possible if the distribution of TEs in the population of genomes follows a Poisson distribution. The main result of this Fig. 1 is that if the distribution of TEs is over-dispersed and the value of *φ* within a given range of values, it suggests that the SE scenario occurs indicating successful invasion of TEs within the genome population.

### 2.3 Numerical methods

We create TEDyS software (**T**ransposable **E**lements **Dy**namic**S**) to simulate the long-term evolution process of TEs through genomes with an individual based model. The Gillespie algorithm (Gillespie, 1976) purpose is to solve a system of differential equations by simulating the occurrences of the events described by the mathematical relations of the equations. Consequently, this algorithm allows the transformation of the model (1) (Flores-Ferrer et al., 2021) to a stochastic individual based model. In the TEDyS software, we combine the Gillespie algorithm of population of genomes with the addition or removal of TEs to each genome to observe and quantify the distribution of TEs over the population of genomes. The mathematical results of (Flores- Ferrer et al., 2021) were obtained supposing specific distributions of TEs and this limitation vanished with the use of the Gillespie algorithm where the entire system is simulated (population of genomes containing TEs). To transform the set of differential equations (1), we list the events occurring:

- Asexual reproduction of a genome: the genome is duplicated with its TEs
- Death of a genome by intra-specific competition or naturally
- Transposition of a TE in its host genome
- Deletion of a TE in its host genome
- Activation of a TE, a silent TE becomes active (the rate is *a*)
- Silencing of a TE, an active TE becomes silent (the rate is *s*)

We firstly implement the Gillespie algorithm from every part of the system (1) but the activation and silencing rate have a difference of order of magnitude of 10^3^ compared to the other values we intend to use. Another reason is that the Co-Extinction scenario starts with an exponential growth of TEs before a simultaneous reduction of host (and consequently, reduction of the number of TEs). Fig. 3A where the exponential increase of TE starts before leading the two populations (TEs and genomes) towards zero. This exponential increase of the number of TEs leads to a drastic slow down of the simulation process because of the rates. To reduce computation time due to activation and silencing of TEs we choose to simplify the model by fixing a proportion of active TEs within the TE population of an individual *p*_*a*_. This implies that, during the Gillespie algorithm, when a TE is added or removed to a genome, the nature of this TE (active or silenced) will be such that the proportion will be as close as possible to this threshold value.

- Asexual reproduction of a genome, it is duplicated with its TEs
- Death of a genome by intra-specific competition or naturally
- Transposition of a TE in its host genome, the TE state (active or silent) is chosen to be closer to the proportion of active TEs in the genome.
- Deletion of a TE in its host genome, a TE is chosen accordingly to its state to be closer to the proportion of active TEs in the genome.

The algorithm runs until a stopping condition is reached; here, a limit of 20,000 time units was chosen to ensure a sufficient number of ‘generations,’ allowing for multiple population renewals on average. Fig. 2 details a state diagram of the algorithm used with a fixed ratio of active TEs in each genome. The next event is randomly chosen with weights depending on the rates of each event (*τ*). Thus, this equation

**Figure 2:**
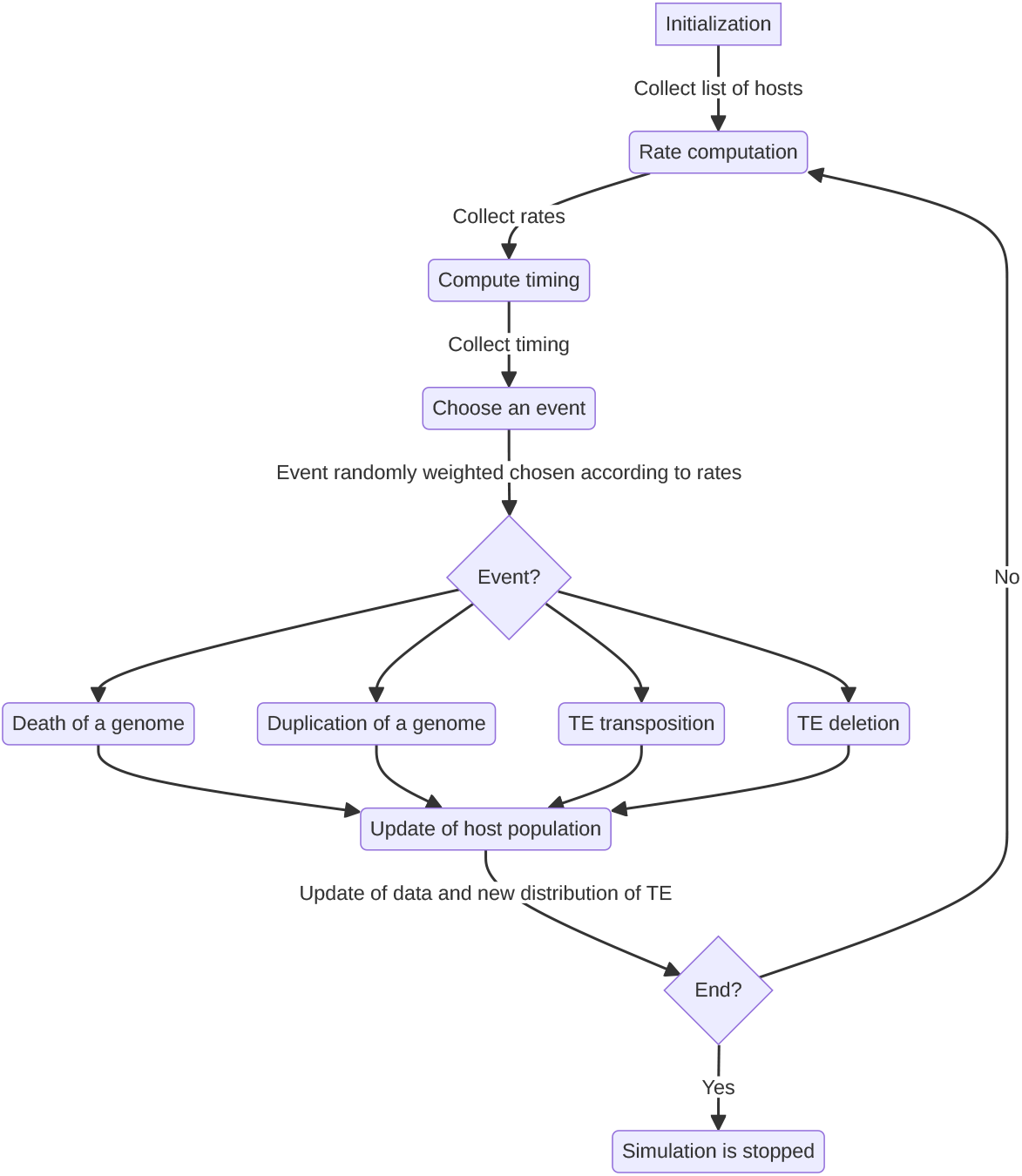
State diagram of the Gillespie algorithm developed to solve the differential equations’ system (1) stochastically. A weight *τ* is computed for each of the four events for each genome of the population. Thus, equation (2) compute the additional time it takes to achieve this event. The simulation is stopped when the maximum time is reached.

**Figure 3:**
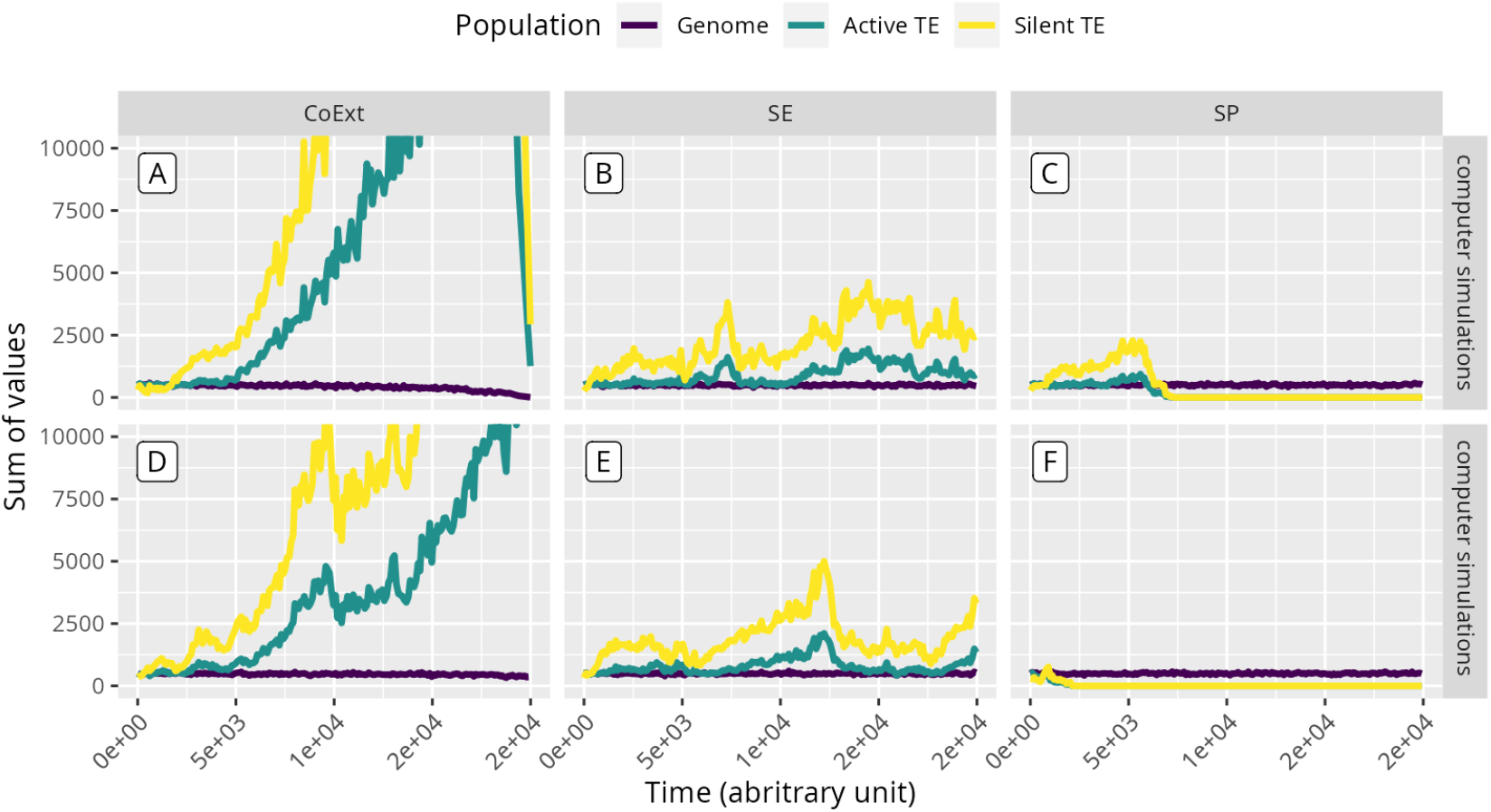
Visualisation of six computer simulations detailing the population of genomes, and the populations of active and silent TEs. These six simulations are part of a series of 30 simulations with the same parameters (Tab. 1). Stochastic processes of TEDyS model can exhibit three scenarios: Co-Extinction for A and D, Stable coexistence Equilibrium for B and E and Selective Purge for C and F. Simulation A show the increase of total number of TEs until they reach a peak that lead the genome population to extinction decreasing drastically the total number of TEs in the population at the end of the simulation. For simulation C and F, the TEs are purged while in simulation B and E, the population of TEs is maintained, indicating a stable coexistence. The host birth value *b*_*h*_ is 0.1, and the host death *d*_*h*_ is 0.05, the intraspecific competition *α* and the deleterious effect *φ* as well as the death rate of TEs *d*_*t*_ have the value of 0.0001, the birth rate of TEs *b*_*t*_ are 0.0015 and the percentage of active TEs *p*_*a*_ is equal to 0.3.

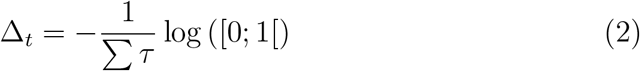

calculates the additional time required for this event to occur.

Consequently, the simulation process chosen is to fix a ratio *p*_*a*_ to force the system to maintain artificially the distribution of TEs to this rate because acceleration methods (Charlebois and Kærn, 2013) cannot be employed as they are based on the distribution of the entities we are seeking to characterise. This approach permits us to simulate the sequence of events that could lead to the three expected scenarios (Co-Ext for co-extinction, SE for stable equilibrium, SP for selective purge). In genetic algorithms, populations are renewed at each generation and selection is based on the fitness of individuals. In contrast, in Gillespie’s algorithm, the population is continually renewed, and selection is based on the calculation of weights for each action in each genome. A genome that accumulates numerous TEs is penalised and the event that leads to its removal from the population is favoured.

In addition, with this computational approach, the distribution of TEs can be recorder in the population of genomes instead of imposing it. The purpose is to monitor the influence of the distribution of TEs in the population of genomes considering two types of initial conditions for the model: the distribution of TEs follows a Poisson distribution or a negative-binomial distribution. These distributions will not persist due to the stochastic properties of the model implemented in TEDyS software.

### 2.4. Computer simulations

**Table 1:** Parameters used for simulation campaign to explore the possibilities of the simulator. As this algorithm is stochastic, we perform 30 simulations for each set of parameters. This lead to a total amount of 87360 simulations to perform and analyse.

| Model parameter | Values | Count |
| --- | --- | --- |
| deleterious effect of TEs $\phi$ | $[10^{-4}, 10^{-3}]$ in steps of $10^{-5}$ | 91 |
| Proportion of active TEs $p_a$ | $[0.3 ; 0.4 ; 0.5 ; 0.6 ; 0.7 ; 0.8 ; 0.9 ; 1]$ | 8 |
| Initial distribution of TEs | Negative binomial or Poisson | 2 |
| Initial average number of TEs | 1 or 5 | 2 |

Total $30 \times 91 \times 8 \times 2 \times 2 = \mathbf{87,360}$

We design an exploratory campaign of simulations to evaluate and analyse the TEDyS software for relevant parameters *φ* the deleterious effect of TEs, *p*_*a*_ the proportion of active TEs and the initial distribution of TEs in the population (Tab. 1). The following values remain constant: *α*_*h*_ = 10^*−*4^, *b*_*h*_ = 0.1, *d*_*h*_ = 0.05, *b*_*t*_ = 0.0015 and *d*_*t*_ = 10^*−*4^. Further to these values, the expected lifespan is 20 time-units, and the simulation maximum time allowed is 20,000 time-units, thus we expect an average of 1,000 generations. Finally, we consider 1 or 5 TEs in average is like the one used in (Le Rouzic et al.,2007) as the average value of these two distributions of TEs to estimate the influence of the initial conditions over the results. Consequently, we performed the 87,360 stochastic simulations for each set of parameters listed in Tab 1.

The output of TEDyS enables the user to monitor regularly the distribution of TEs in the population of genomes. This first simulation campaign has been completed to a second one with a choice of parameters leading to the three scenarios identified (Co-Ext, SE and SP) while exporting the distributions more frequently to understand their dynamics. Same parameters as previously except for *p*_*a*_ and *φ*:

1. CoExt parameters: *φ* = 10^*−*4^ and *p*_*a*_ = 1
2. SE parameters: *φ* = 5.10^*−*4^ and *p*_*a*_ = 0.3
3. SP parameters: *φ* = 5.10^*−*3^, *p*_*a*_ = 1

### 2.5. Post-treatment of the simulations

The Fig. 3 details potential scenarios TEDyS software can produce. Fig. 3D is specifically labelled `CoExt_ongoing` because the exponential growth has not yet reached its peak before the population collapse, unlike in Fig. 3A. However, as there are still TE and hosts in simulation A of Fig. 3, as well, such a simulation is also labelled `CoExt_ongoing.` This Fig. 3 present various scenarios while the parameters are the same, underlying the stochastic aspect of our simulator.

After a post-treatment of these simulations output, we are able to estimate the parameters’ adjustment to the Poisson distribution and the negative-binomial distribution of TEs (the latter is referred as the over-dispersed distribution in Fig. 1). The following criteria are used to discriminate the simulations and attribute a scenario:

`CoExt_end` The simulation results in no more TE or hosts remaining before the maximum allotted time (20,000 time-units) is reached.

`CoExt_ongoing` The simulation is currently reducing the average number of hosts, while the average number of TEs is increasing following an exponential trend.

`CoExt_start` The host population remains constant around the equilibrium value. The increase in the average number of TEs follows an exponential law.

`SE_start` The host population remains constant around the equilibrium value. The increase in the average number of TEs follows a logistic growth pattern.

`SE` The host population remains constant around the equilibrium value, and the TE population remains relatively constant.

`SP_start` The average TEs population decreases to a value below 1, indicating an upcoming purge of TEs.

`SP` There are no more TEs in the host population.

To differentiate between the `CoExt_start` and `SE_start` scenarios, we fitted the average value of TE in individuals to both an exponential and logistic growth model, then selected the scenario based on a visual comparison of the two fits in addition of the statistical values characterizing the fits and their Akaike weights (Akaike, 1974) (see Portet (2020) for a recent review on model selection using this criterion).

Beyond population dynamics, the TEDyS software monitors the distributions of TEs over time by computing their mean and variance values. Computing the variance and the variance-to-mean ratio provides insight into the dispersion of the distribution and allows assessment of whether the data follow a Poisson-like process or exhibit over- or under-dispersion. During post-treatment, the distribution of TEs is also adjusted to two distributions Negative-Binomial and Poisson. The Poisson distribution is adjusted with the mean value *λ* and reciprocally the negative binomial distribution is adjusted via two parameters *µ* (the mean value) and *k* the aggregation parameter. Smooth regressions over time of parameters *k* are done using loess (local polynomial regression fitting) function of R package `stats.` To complement this post-treatment, Akaike weights are also used to quantify the best fitting distribution of TEs: negative binomial distribution (weight *w*_nb_) is compared to Poisson distribution (weight *w*_*P*_). In order to facilitate the monitoring of progression, additional visualisations of the value distributions are provided at 10% intervals, offering a clearer representation of their temporal evolution. Recurrence plots, as introduced and further developed by Eckmann et al. (1987) and Kurths (Marwan et al., 2007), provide a powerful tool for revealing hidden patterns and nonlinear structures in the dynamics of complex systems. Density histograms display the distribution of a dataset by showing the relative frequency of observations across intervals, normalized to represent probability density. Statistical analysis were carried out using R 4.5.0 (R Core Team, 2021) with tidyverse (Wickham et al., 2019), MASS (Venables and Ripley, 2002) and vroom (Hester et al., 2024) libraries.

**Table 2:** Distribution of scenario over the 87,360 simulations listed in Tab. 1. The most challenging cases to distinguish represent only less than 5% of the total simulations conducted.

| CoExt<br>39.3% |  |  | SE<br>5.5% |  | SP<br>55.2% |  |
| --- | --- | --- | --- | --- | --- | --- |
| CoExt_end<br>35% | CoExt_ongoing<br>3.2% | CoExt_start<br>1.1% | SE_start<br>0.3% | SE<br>5.2% | SP_start<br>0.2% | SP<br>55% |

## 3. Results

### 3.1. Emergence of Stable Equilibrium

The purpose of TEDyS model is to simulate an invasion of TE in genomes for asexual reproduction within the modelling framework of host-parasite interactions where TEs are considered as deleterious. As demonstrated analytically (Flores-Ferrer et al., 2021) the distribution of TEs within the population of genomes has an impact on the scenarios this system can produce. As the deleterious effect of the TEs is fixed all along the simulation we studied the increase of parameter *φ* dedicated to the deleterious effect of TEs to a genome.

#### Deleterious effect and scenarios

In appendix A, Fig. A.12 shows the simulation campaign as a whole (post-treatment of the 87,360 simulations) and gives an idea of the scenarios that can be observed according to the different parameters selected. The overall results are grouped, and the scenarios counted to have a glimpse of their relative importance (Tab. 2). Focusing on the main scenarios (CoExt, SE and SP), Fig. 4 summarise the result for *p*_*a*_ ≥ 0.5. The simulations are done for 20,000 time-units in the Gillespie algorithm corresponding to an average value of 1,000 generations according to parameters chosen (2.4). For most of the plots of Fig. 4, the transition from co-extinction to selective purge is clearly reproduced when *φ* is increased. The initial conditions chosen corresponds to an unrealistic state, but they are only used here to analyse long-term evolution of TEs as it has been done in (Le Rouzic et al., 2007). The initial condition of the second column (1 TE in average with a Poisson distribution) mainly lead to selective purge scenarios. As expected by theoretical results, Fig. 4 underline that it exists a range of values of *φ* for witch a majority of simulations maintain TEs coexistence in the population of genomes when the parameter *p*_*a*_ is equal to 0.3 (30% of active TEs in the genome in average). Increasing the average proportion of active TEs (*p*_*a*_) in genomes reduces the number of occurrences of stable equilibrium scenario.

**Figure 4:**
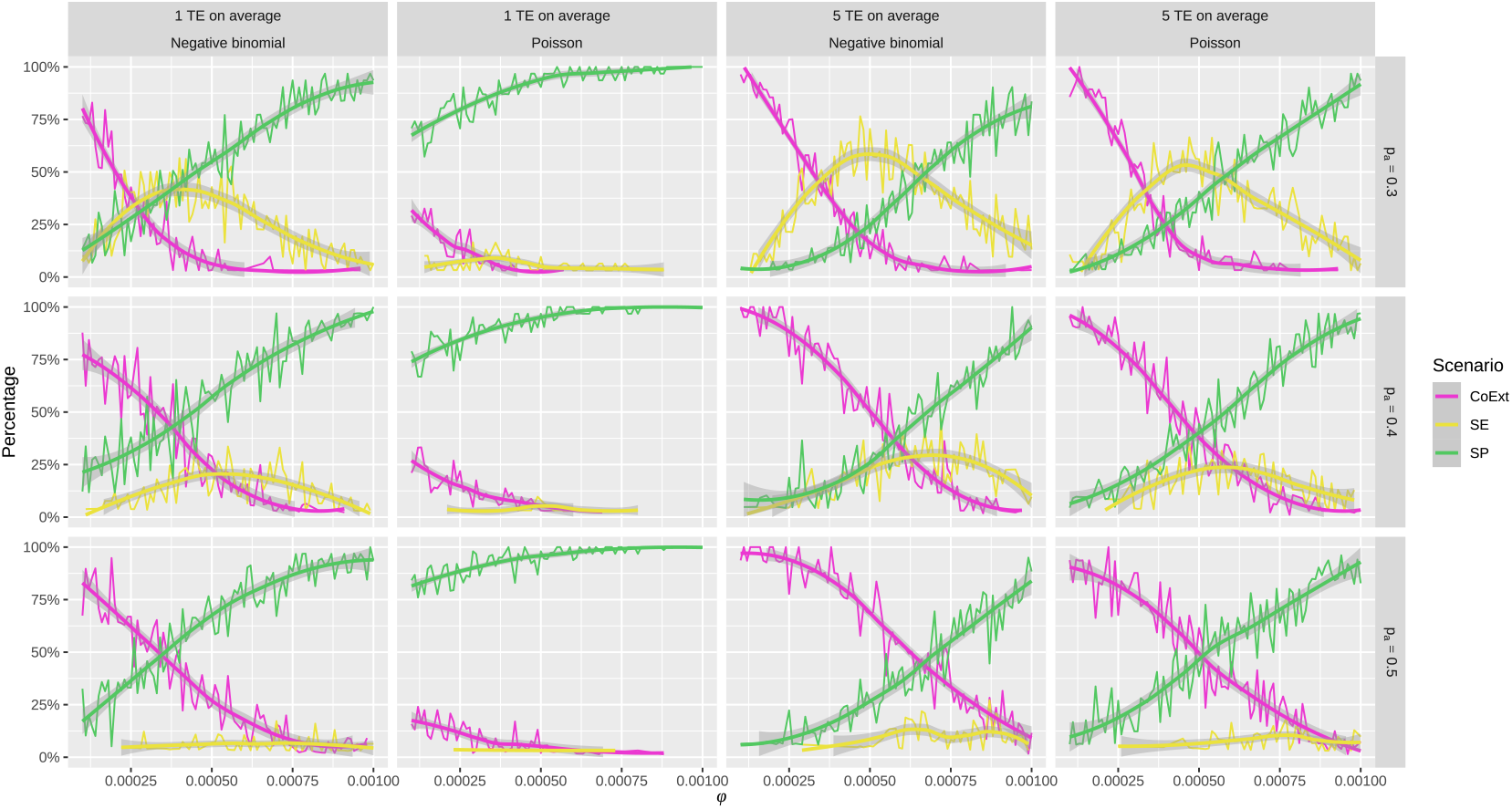
Percentage of scenario for 30 simulations depending on *φ* and *p*_*a*_. Only simulations with a value of *p*_*a*_ greater or equal than 0.5 are represented. Fig. A.12 gives the full representation of the computer simulations. Scenarios are split into three categories (CoExt, SE, and SP). The ratios underline the predominant scenario as *φ* increases detailing the transition from CoExt to SP. The plot represents smooth regression using loess (local polynomial regression fitting) function of R package `stats` over 30 replicates.

For values of *p*_*a*_ higher than the threshold of 0.5, there are rare occurrences of the SE scenario (Fig. A.12). Conversely, for values below this threshold, co-extinction scenarios are predominant when the value of *φ* is close to 10^*−*4^, whereas at the opposite end of the *φ* range, selective purge scenarios dominate. These initial results seem to support the possible emergence of SE scenarios in the middle of this *φ* interval, as predicted by the analytical results (Fig. 1 (Flores-Ferrer et al., 2021)).

#### Aggregation parameter k

As stated in (Flores-Ferrer et al., 2021) the distribution of TEs has a significant impact on the expected scenarios the system could reach (Fig. 1). When the aggregation parameter increases, this indicates a lack of aggregation corresponding to Poisson distribution. This induces a removal of points of Fig. 5A to obtain Fig. 5B where only simulations following a negative distribution are conserved according to Akaike criterion testing Poisson versus negative binomial distribution.

**Figure 5:**
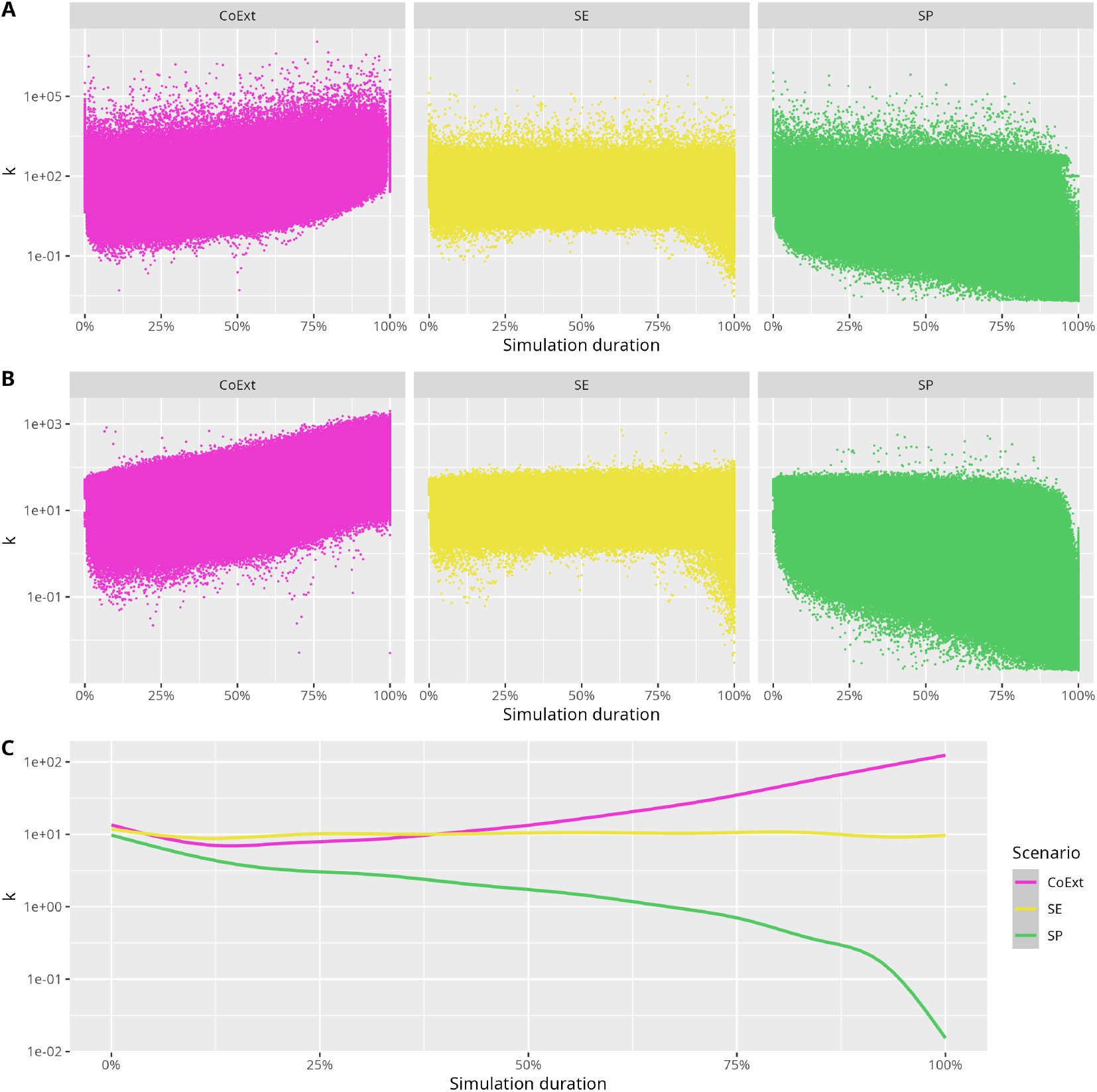
Average value of *k* during a simulation. The simulation duration is rescaled to percentage to compare SE simulations with purge where TEs disappear fast, CoExt simulations where both populations collapse and SE simulations where TEs persist. A corresponds to all values of *k* fitted and B, only the values when the distribution follows a Negative Binomial distribution according to Akaike weights (*w*_nb_ *>* 0.5). C. The plot represents a smooth regression of the values of B. Selective purge scenarios (SP) tends to an aggregation parameter *k* toward 0 because of the progressive removal of the TEs. Co-extinction scenarios (CoExt) that reached the end by crashing both populations (Host and TEs) indicate that the aggregation parameter *k* exponentially increase during the simulation. Stable equilibrium scenarios (SE) have an aggregation parameter that remains constant with an average value of *k* = 10.

Thus, the aggregation parameter *k* is used to evaluate the impact of the structure of the distribution to the scenario. Fig. 5 indicates that the initial conditions do not have an impact on the structure of the aggregation related to the scenarios. For the co-extinction scenario the trajectories indicate a quasi-exponential growth of *k* during the simulation. Reciprocally, for stable equilibrium a plateau is maintained throughout the simulation. These results (Fig. 5) suggest that the aggregation of TEs occurs progressively until coextinction, whereas, in other scenarios, diversity is maintained from the very first iterations of the model. Consequently, diversity is preserved up to a point where accumulation leads to co-extinction or, conversely, the spread of the distribution is so extensive that all individuals are different, triggering selective purging. It follows that the parameters and the model are such that the system could sustain with a diverse distribution of individuals for a long time, with varying numbers of TEs, allowing the TEs population to persist within the genome population. Analysis of Fig. 5C provides an overview of the parameters when the distribution of TEs potentially follows a negative binomial distribution more closely. The shifts from this distribution to Poisson distribution are examined in a later section.

### 3.2. Persistence of TEs in Stable Equilibrium scenarios

This Fig. 6 demonstrates the feasibility to our model to maintain for a long time (more than an average of 10,000 generations) the TEs in the population of genomes. The numerical results of Fig. 7 confirm the theoretical approach by showing the longest seventeen simulations for a value of *φ* closed to the threshold *r*_*T*_ = *p*_*a*_(*b*_*T*_ − *d*_*T*_) − (1 − *p*_*a*_)*d*_*T*_ = 3.5*e*^*−*4^ for these specifics simulations (Fig. 1). As expected, moving away from this value of *r*_*T*_, the duration of the simulations decreases. Fig. 7 provides the average number of TEs evolution for the seventeen select simulations demonstrating the robustness of the model to maintain TEs in the population of genomes for a long time (an average of 10,000 generation). The parameters of the system maintain an average number of TEs for an average value of 18,000 generations. Even if all simulations end by one of the two scenarios inducing TEs extinction (CoExt or SP), the average number of generations that persist is enough to demonstrate that deleterious TEs can maintain without any change of their parameters or other mechanism.

**Figure 6:**
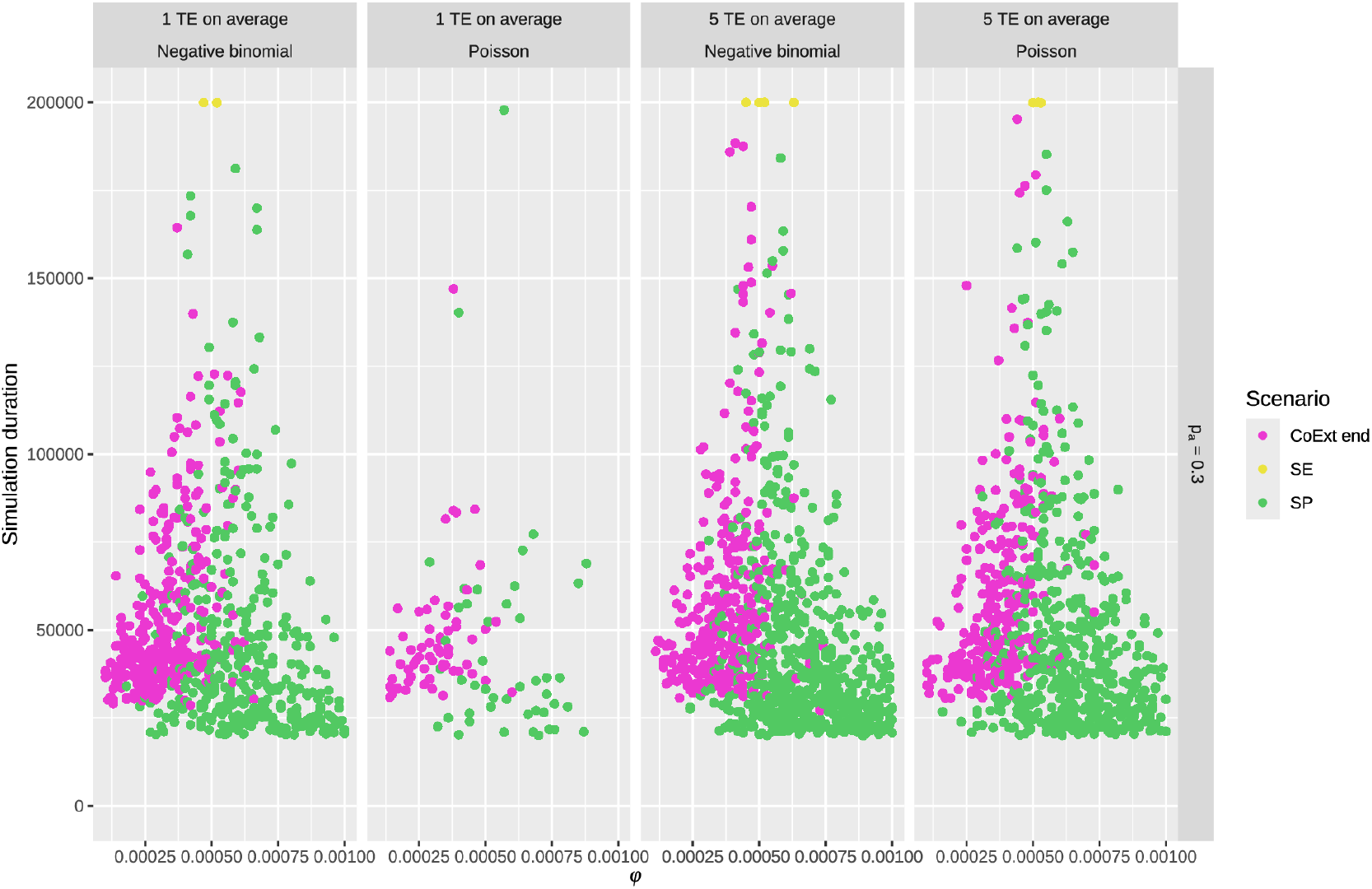
Selected stable equilibrium scenarios of Fig. 4 for ten times longer simulation time. The 17 longest simulations have a parameter value close to *φ* = 5*e*^*−*4^ that is the theoretical value *r*_*T*_ = 3.5*e*^*−*4^. These plots indicate that the initial conditions do not impact the results allowing long simulations for each. 200,000 time-units correspond to an average number of 10,000 generations.

**Figure 7:**
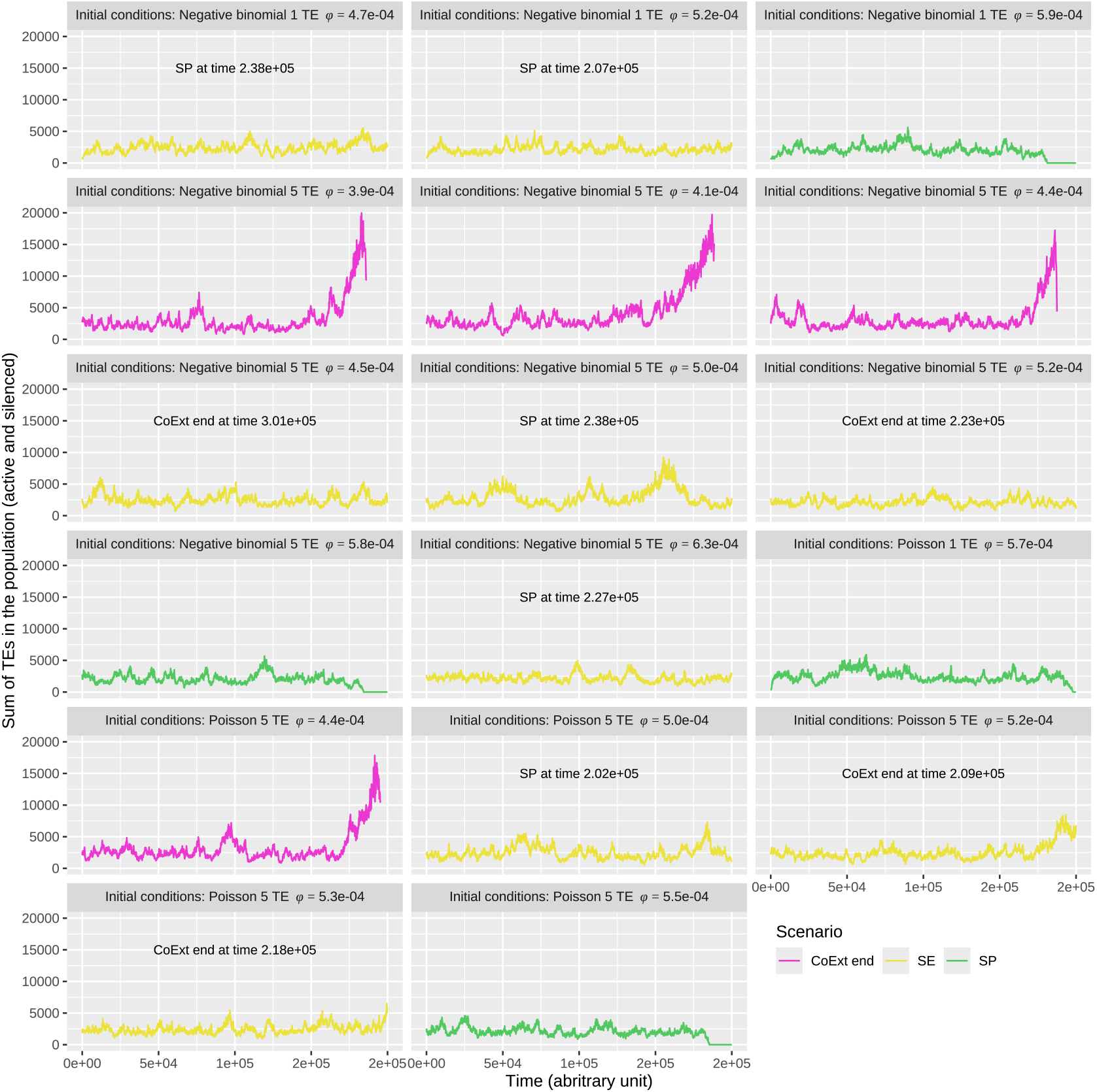
17 simulations maintaining TEs beyond 180,000 time-units. 9 of them go beyond the threshold of 200,000 time-units and the maximum time-units reached is indicated on the plot as well as the scenario. One of them runs to an average value of more than 30,000 generations. *p*_*a*_ = 0.3 for these simulations.

### 3.3. TEs dynamics and distributions

For a better understanding of dynamics, we have chosen to keep only the portions of the simulations that have TEs and to group them into ten percent portions. Three figures show the evolution of Akaike’s weight *w*_nb_ (distribution of a negative binomial distribution versus a Poisson distribution) according to the simulation portion and the scenario (Figs. B.13, 8 and 9). Fig. B.13 shows the boxplots which indicates that the distribution of weights is the same throughout the SE simulations, suggesting a certain stability in this case. Variations over time are observed for the other scenarios as illustrated by Fig. 8 with density histograms of the Akaike weight *w*_nb_. For CoExt, towards the middle of the simulations leading up to it, almost all the simulations follow a negative binomial distribution before having a mixture of Poisson and negative binomial at the end when co-extinction occurs. For SE, the distribution is the same all along the simulations indicating that the dynamics is maintained as well as TEs. Finally, for the selective purging scenarios, the structure of the distribution mostly follows a Poisson distribution, except for a few simulations at the end (last ten percents).

**Figure 8:**
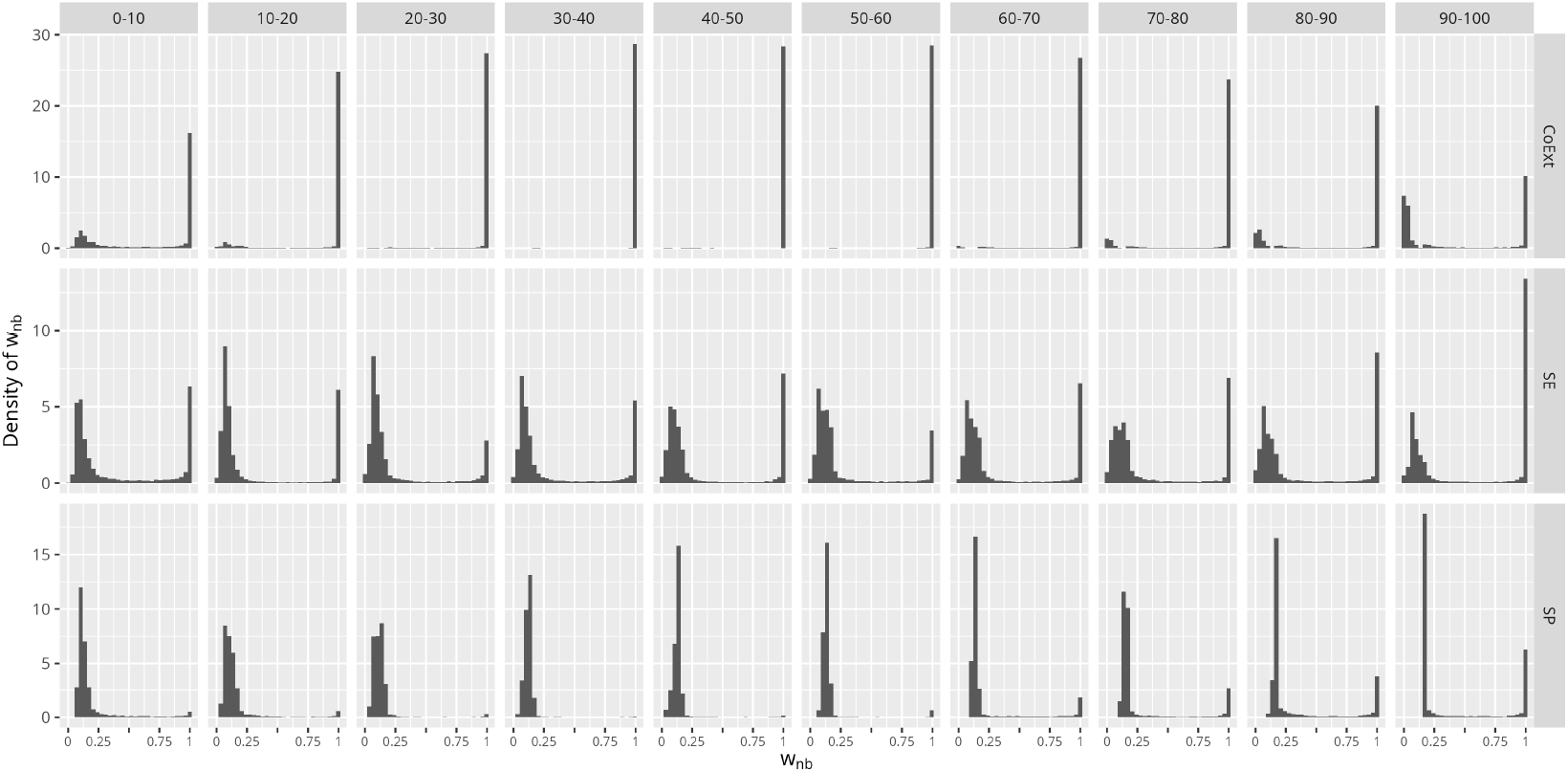
Normalised distribution of Akaike weights of the negative binomial distribution compared to a Poisson distribution over block of 10% of simulation duration. Simulations are split into ten slices to have a glimpse of the evolution of the repartition of Akaike weights of the negative binomial distribution (*w*_nb_) with density histograms.

The recurrence diagrams (Fig. 9) show whether there are any patterns in the distribution of the successive values of the Akaike weights. We note that in the middle of the simulations leading to the removal of the TEs (SP or CoExt), there are no longer any transitions between the two types of distributions - it’s one or the other. Conversely, throughout the SE simulation, these transitions between the two distributions are frequent and maintained. This is also the case at the end of the SP and CoExt simulations, but these are the last moments in the simulations and the number of TEs, or individuals involved is low. On the other hand, these transitions are not visible in the middle of these simulations, which shows a significant portion where the distributions followed a distribution rather close to a negative binomial distribution for CoExt and Poisson distribution for SP.

**Figure 9:**
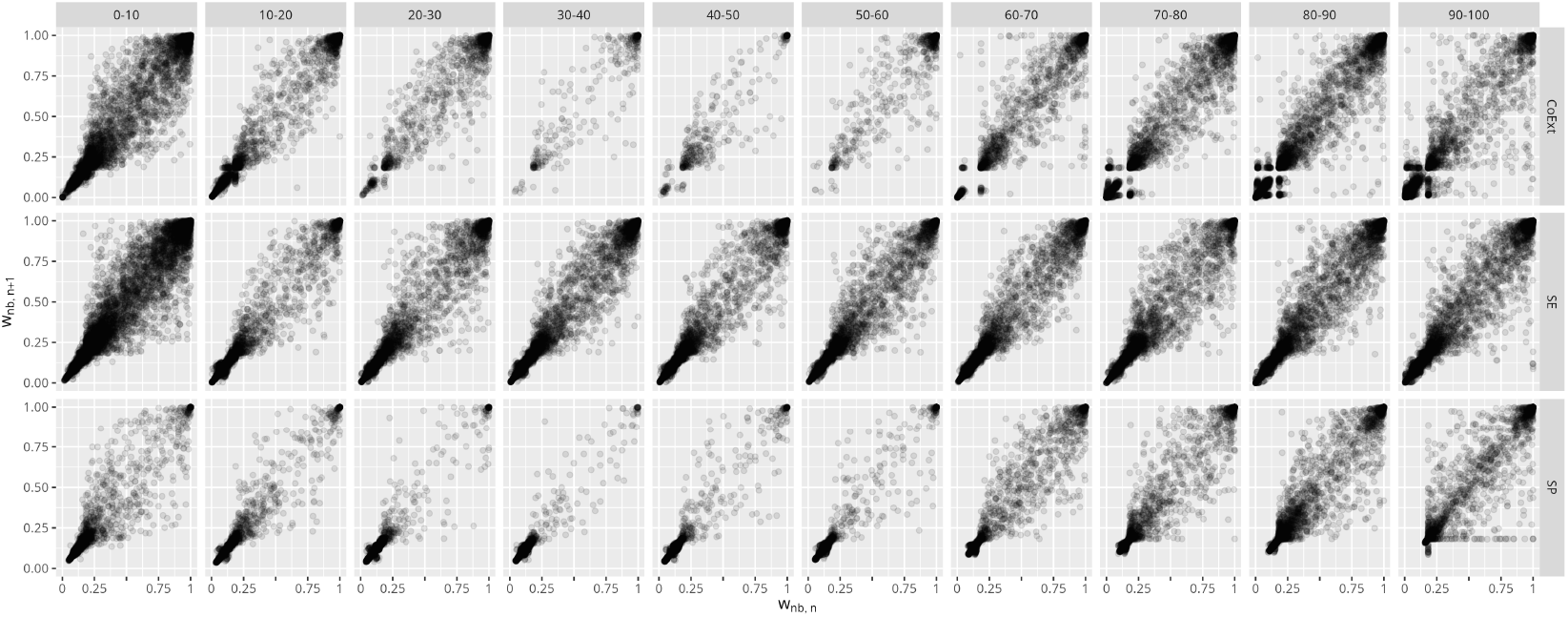
Recurrence map to have a glimpse of successive values of histograms Fig. 8. Simulations are split into ten slices to have a glimpse of the dynamic evolution of Akaike weights of the negative binomial distribution (*w*_nb_). For CoExt and SP scenarios, shape evolution reflects changes in dynamics. For SE scenarios, stable diagram shape across segments, indicating process continuity.

In the rest of the results, segmented according to scenarios, the Fig. 10 represent the computed values for a set of 10 simulations over 30 performed. These are the evolution of series of parameters and measurements to clarify the dynamics of TEs during the simulations. Fig. 10A displays that the population of TEs exponentially increases until the end of the simulation while the host population decreases causing the co-extinction of both populations. The distribution characterised by Akaike weights is mostly negative binomial until a point where there are so many TEs in few genomes that the distribution of TEs follows a Poisson distribution more than a negative binomial. Bursts of TEs are observed continuously, and they lead the distribution of TEs to a negative binomial distribution. There are so many TEs in the genome that the selection process can no longer take place and so the accumulation of TEs occurs by maintaining a negative binomial distribution. The accumulation is such that, after a while, the population falls and the number of individuals decreases, so that, with so few individuals, the best-fitting distribution is a Poisson distribution.

**Figure 10:**
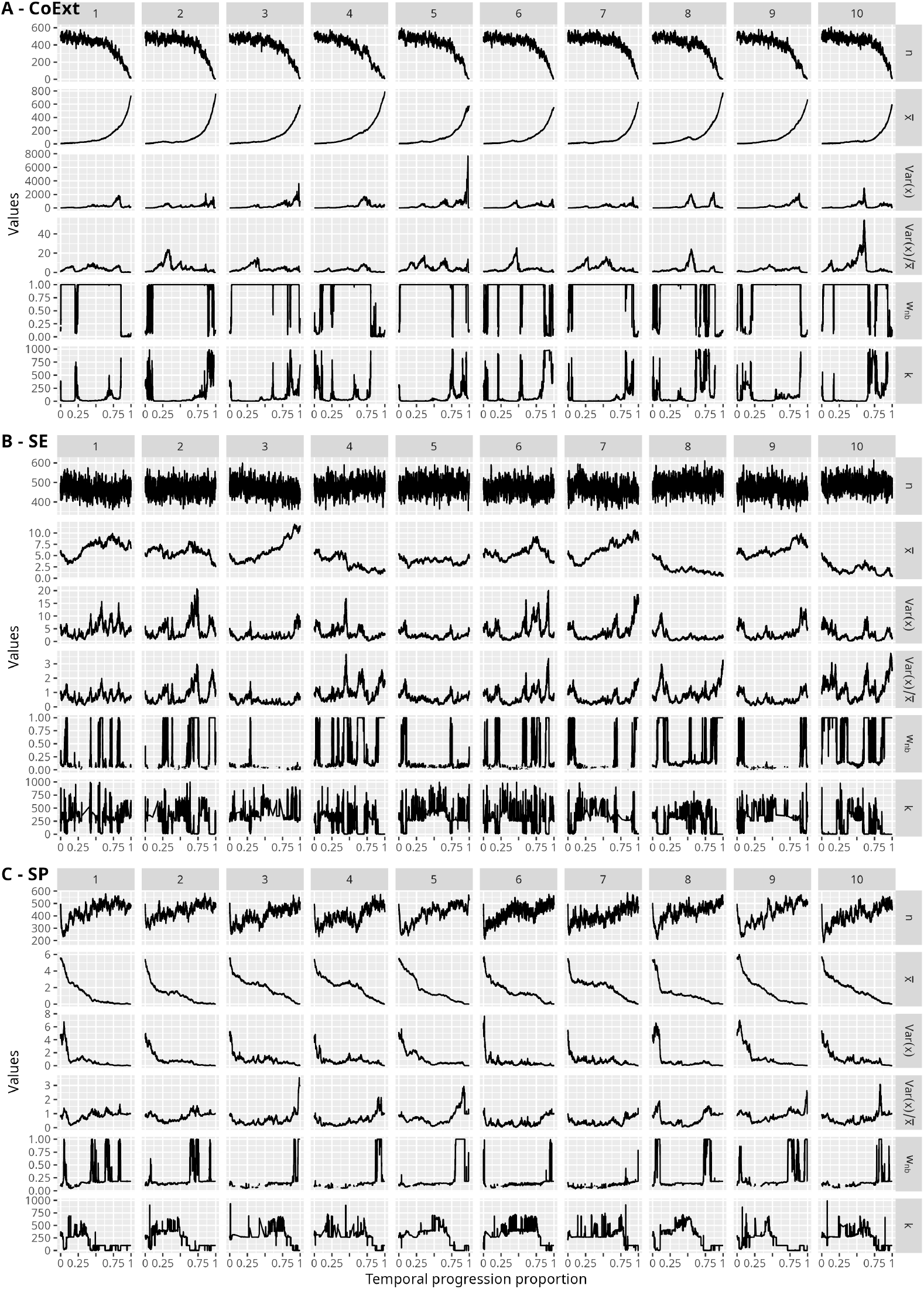
Evolution of characteristics during (A) Co-Extinction simulations, (B) Stable Equilibrium and (C) - Selective Purge scenarios. Values are *n* the number of genomes, 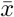 the average number 1of TEs per individual over the population, Var(*x*) the variance of the distribution of TEs per individual, Var(*x*)/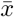 the variance-to-mean ratio, also referred to as the index of dispersion, *w*_nb_ the Akaike weight of the negative binomial distribution compared to a Poisson distribution and *k* the aggregation parameter when the distribution if fitted with a negative binomial distribution.

Fig. 10C indicates that the population of genomes is maintained to an average of 500 individuals after an initial drop due to the deleterious effect of TEs. At the end of the simulation, TEs are purged. This process is either fast or few TEs can persist before being purged. This leads to a distinct adjustment to distribution of TEs at the end. In the fast case, the distribution of TEs follows a negative binomial distribution and, respectively, when few TEs are maintained in the population, the distribution is Poisson. It is important to note that, due to stochasticity, it might happen that few TEs (less than 1 TE in average in the population) could lead to a reinvasion of TEs within the genomes where the average value is 5 TEs. However, these results in a population that is overly diverse and unable (given the processes of the differential equation system) maintaining a subpopulation with a few numbers of TEs, ultimately leading to their purging.

Considering SE scenarios, TEs are maintained all along the simulation within host population of genomes (Fig. 10B). There are multiple shifts in the distributions of TEs: the distribution follows a Negative Binomial distribution for an average period of 35 time-units. When moving from a Poisson to a Negative binomial distribution, it mostly (in 70% of the cases) corresponds to an increase of the average number of TE and of the variance of the distribution (Figs. 10 and 11). Reciprocally, from a Negative Binomial distribution to a Poisson distribution, it corresponds to a decrease of the average number of TEs in the same proportion (70% of the cases). During the simulations, a burst of TEs is observed in 70% of the case when the population become less aggregated. This burst is under control of the model where the parameter of the deleterious effect of TEs is set to prevent exponential growth leading to co-extinction. By extending the simulations for a long time, the random choice sequences will eventually lead these simulations towards one of the other scenarios. These parameters ensure that most simulations are maintained over a long period of time.

**Figure 11:**
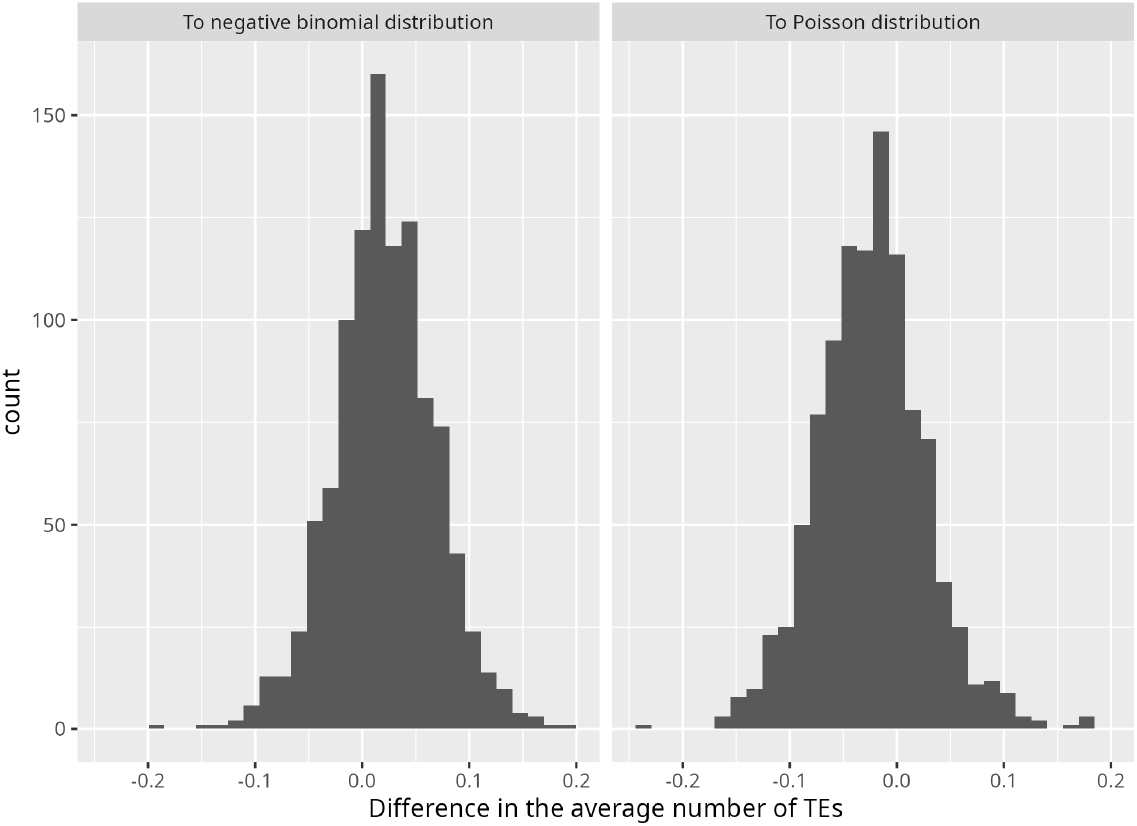
For SE scenarios, histograms of the differences in the average number of TEs depending on the distribution towards which the simulation is evolving. These transitions from one distribution to another are detected using Akaike weights. 1,050 observations for histogram entitled “To negative binomial distribution” where 68% of values are positive and 1,040 observations for histogram entitled “To Poisson distribution” where 70% of values are negative.

## 4. Discussion

### 4.1. Diversity of TE population

The results indicate that, beyond the quantity of TEs accumulated in the genomes of a population, the diversity of situations with a relative aggregation of TEs enables the long-term maintenance of the TE population in genomes under selective pressure due to the deleterious effects of TEs. Indeed, as shown in Fig. 5, there are scenarios where TEs accumulate with diversity in co-extinction scenario, leading to a progressively exponential trajectory in the evolution of the aggregation parameter *k*. Consequently, the entire genome population gradually becomes trapped in TE accumulation leading to a co-extinction scenario. In contrast, for stable equilibrium scenarios, the evolution curve of the parameter *k* quickly reaches an equilibrium value, indicating that diversity is rapidly established and maintained. For selective purge scenario, the diversity is weaker with a downward trend for a long period before declining. This decline signifies that, gradually, the disparity in the genome population becomes too high, and all genomes are subjected to selection, with individuals harboring fewer deleterious TEs being eliminated, leading to a purge. The TEDyS simulations reveal an interesting mechanism where, within the genome population, multiple aggregates form. If one of these aggregates begins to accumulate too many TEs, it is eliminated by selection pressure (Supplementary material Video 1). For co-extinction scenarios, burst of TEs implies that the distribution of TEs follows a negative binomial distribution for long period compared to shorter period where this type of distribution is followed for stable equilibrium scenarios. It implies that for co-extinction scenarios, the overdispersed distribution is maintained having a diversity in the population. The deleterious effect modelled with *φ* is inefficient to prevent accumulation. Reciprocally, for stable equilibrium scenarios, the distribution of TEs indicating burst of TEs are short. It implies that the population is diverse temporarily, but the selection mechanism does its task properly by removing genomes with the higher number of TEs in 70% of the cases (Fig. 11). The stochasticity of the model induced that, sometimes, the purge of individual with more TEs is not done successively finally leading to co-extinction.

This diversity within the population combined with selection allows TEs to persist and creates a window for their co-existence. The equilibrium of parameters balances diversity generation of new genomes with an appropriate deleterious effect to maintain the TEs in the population of hosts. We further demonstrated that, even though these simulations maintained TEs for a long time (few reaching an average of 20,000 generations), the outcome is either co-extinction or the purge. Thus, diversity within the genome population with burst of TEs managed by selection allows TEs to persist for an extended period until the stochastic processes of the model drive either selective purge or co-extinction. Taking into account a potential distribution other than Poisson for asexual reproduction in our model shows that the expected results are different from those hoped (Hickey, 1982; Charlesworth and Charlesworth, 1983) for and that this may also have an impact on sexual reproduction, where Poisson’s law prevail (Dolgin and Charlesworth, 2006; Roze, 2023).

### 4.2. Silencing effect

The proportion of active TEs in the system influence the number of simulations maintaining the stable equilibrium for long period (Fig. 4). Consequently, increasing this proportion of silenced TEs can be seen as a tool to regulate the genome’s defence against TEs invasion. The higher the activity of TEs, the shorter the period with TEs in the population of genomes. Silencing mechanisms allow for a larger window and the prolonged maintenance of TEs within the genome population. This long-time interval is sufficient for a mutation or an environmental change to subsequently enable the persistence of this TEs population within the genome.

### 4.3. Model approximations

More than half of the simulations depict SP scenarios (Tab. 2), with a particularly large proportion of these scenarios occurring in the second column of Fig. 4. We suppose that the initial Poisson distribution with an average value of one TE often leads the host to have only one TE. In the software, a unique TE must be active to initiate the process or transposition and deletion in the genome even if the percentage of active TE is below 1 with the rules used for initial conditions with at least one TE in average leading to a particular initial Poisson distribution.

However, it turns out that this theoretical prediction held true regardless of whether a silencing mechanism was present. As shown in Fig. A.12, this is not the case, since the absence of silencing should correspond to *p*_*a*_ = 1, and in this situation, less than 0.02% of the simulations resulted in a stable equilibrium scenario for a maximum of 20,000 time-units. This might be due to the choice of fixing the proportion of TEs active by the model instead of allowing Gillespie algorithm managing it with the consequences already exposed in terms of computation time. Gradually decreasing the proportion of active TEs in the genome leads to a steady rise in the percentage of simulations reaching the equilibrium state where both populations co-exist for an average of 1,000 generations.

### 4.4. Concluding remarks

The results clearly demonstrate the ability of TEDyS software to mostly reproduce theoretical result (Flores-Ferrer et al., 2021) by demonstrating that it is stochastically possible to maintain active TEs in asexual genomes’ population in the framework of host-parasite interactions where TEs are deleterious. Although TEs do not solely have deleterious effects, as implemented in TEDyS, this software highlights the value of a non-overlapping population-based model for understanding the dynamics of these elements. Diversity within a genome population is fundamental to maintaining TEs within genomes in the long-term. This property combined with an appropriate selection pressure ensures persistence of TEs within the population of genomes. Furthermore, the silencing mechanism plays a crucial role in maintaining a certain quantity of TEs in genomes, limiting the spread of TEs, and preventing them from dispersing excessively. This avoids the failure to aggregate subpopulations of genomes where TEs could persist. This mechanism could also be viewed as a complementary to genetic population models including the trap model of piRNA clusters (Tomar et al., 2023) by allowing enough generation in the simulation process to permit a defensive strategy against TEs as well as environmental adaptation. Tomar et al. (2023) have shown that it is possible to maintain a population of TEs for a long-term only if TE cluster insertions are deleterious as well. This does not contradict our results, since only the quantity of TEs is considered. These piRNA clusters has also been implemented with agent based modelling, and only size and architecture of piRNAs clusters matters (Kofler, 2019): the genome size, the recombination rate and the population size have little influence on the results. The main advantage of TEDyS model is that it does not require any change of TE properties during simulation process (activation or transposition rates) to enable long-term persistence of TEs. In an asexual framework, by having TEs properties set with fixed parameters along the simulation process, deleterious TEs do not prevent a long-term persistence of TEs in genomes. This theoretical model not only accounts for the longterm maintenance of TEs within a host population, but also demonstrates that an increase in the proportion of silenced TEs constitutes a mechanism that promotes such persistence, in agreement with the biological results observed in bdelloid rotifers (Nowell et al., 2021). This TEDyS agent-based model still has many parameters to study to understand the influence of the equilibrium values set by the parameters on the dynamics of TEs.

## Acknowledgments

This study was supported by the MITI project: “Invasions biologiques et ressources hétérogènes : de la modélisation hybride à la gestion EcoBio-Sociale” founded by the Mission pour les Initatives Transverse et In-terdisciplinaires of the CNRS. This study is set within the framework of the “Laboratoires d’Excellences (LABEX)” TULIP (ANR-10-LABX-41) and of the “École Universitaire de Recherche (EUR)” TULIP-GS (ANR-18-EURE-0019). Martin Rosalie thanks Théophile Yvars, Marc-Alexandre Espiaut and Sara Ibrahim for their help with the TEDyS software during their training as part of the Theoretical Biology team at the LGDP.

## Author contributions

Conceptualization: SG. Methodology: MR, SG. Investigation: MR, SG. Visualization: MR. Funding acquisition: SG. Supervision: MR, SG. Writing-original draft: MR. Writing-review & editing: All authors.

## Data and materials availability

All data needed to evaluate the conclusions in the paper are present in the main text and/or the Supplementary Materials. The TEDyS software is openly available on GitHub at https://github.com/mart1rosalie/TEDyS. The R scripts used to generate the plots are deposited in Zenodo https://doi.org/10.5281/zenodo.17218660

## Appendix A. Post-treatment of the simulation

**Figure A.12:**
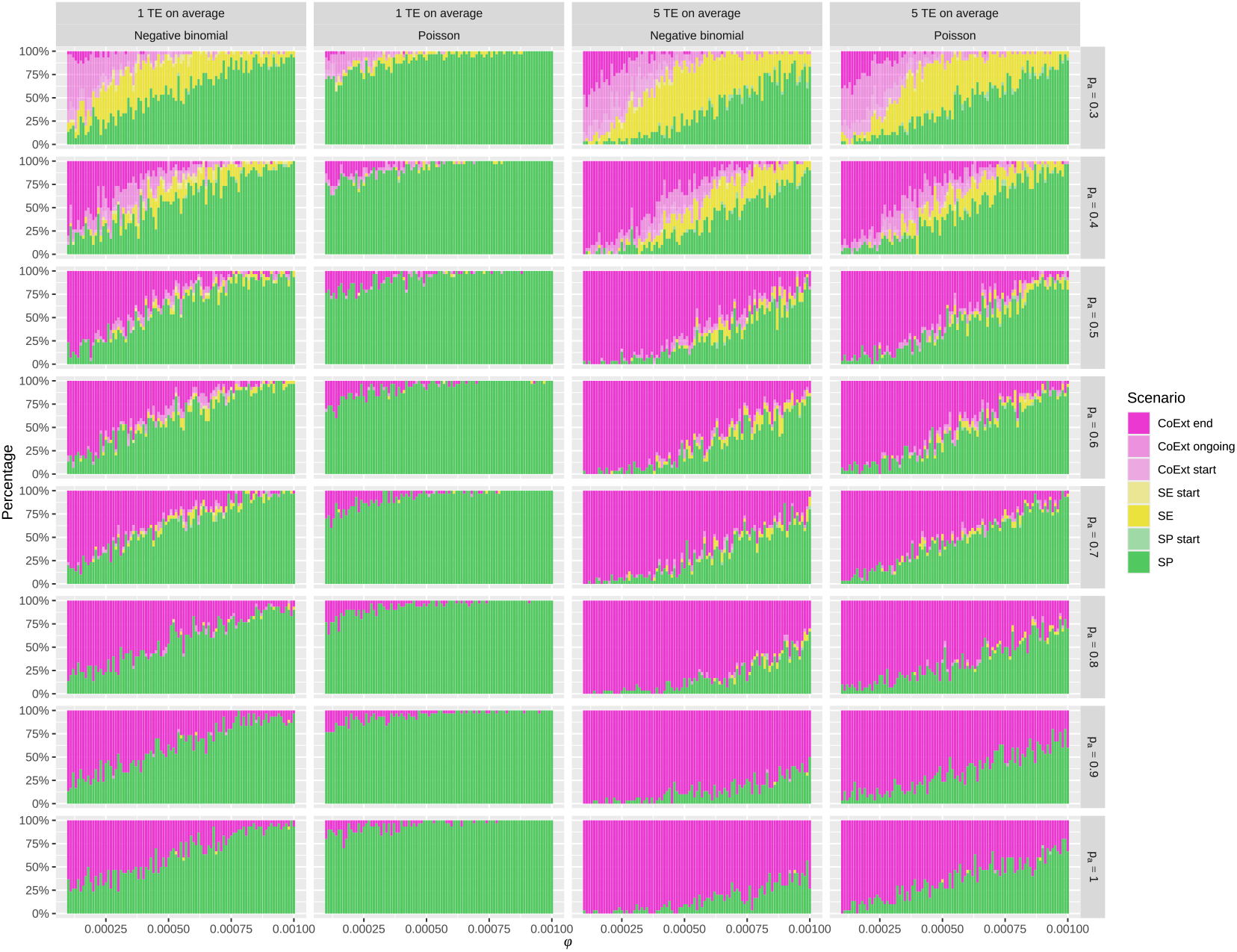
Raw output of the simulation campaign of the 87,360 simulations. The detailed proportion of scenarios of the simulations are displayed across 32 figures arranged in 8 rows and 4 columns. Each figure represents the percentage distribution of the seven scenarios on the y-axis for 30 simulations, and on the x-axis, the variation of the parameter *φ*, which represents the deleterious effect of TEs. The 32 figures are organized into height rows, with the proportion *p*_*a*_ decreasing from 1 to 0.3 from bottom to top, and into four columns with different combinations of initial conditions (Tab. 1). The main categories of scenario are CoExt for Co-Extinction, SE for Stable Coexistence Equilibrium, and SP for Selective Purge and their color correspond to the color of analytical results of Fig. 1.

## Appendix B. Boxplots of Akaike weights

**Figure B.13:**
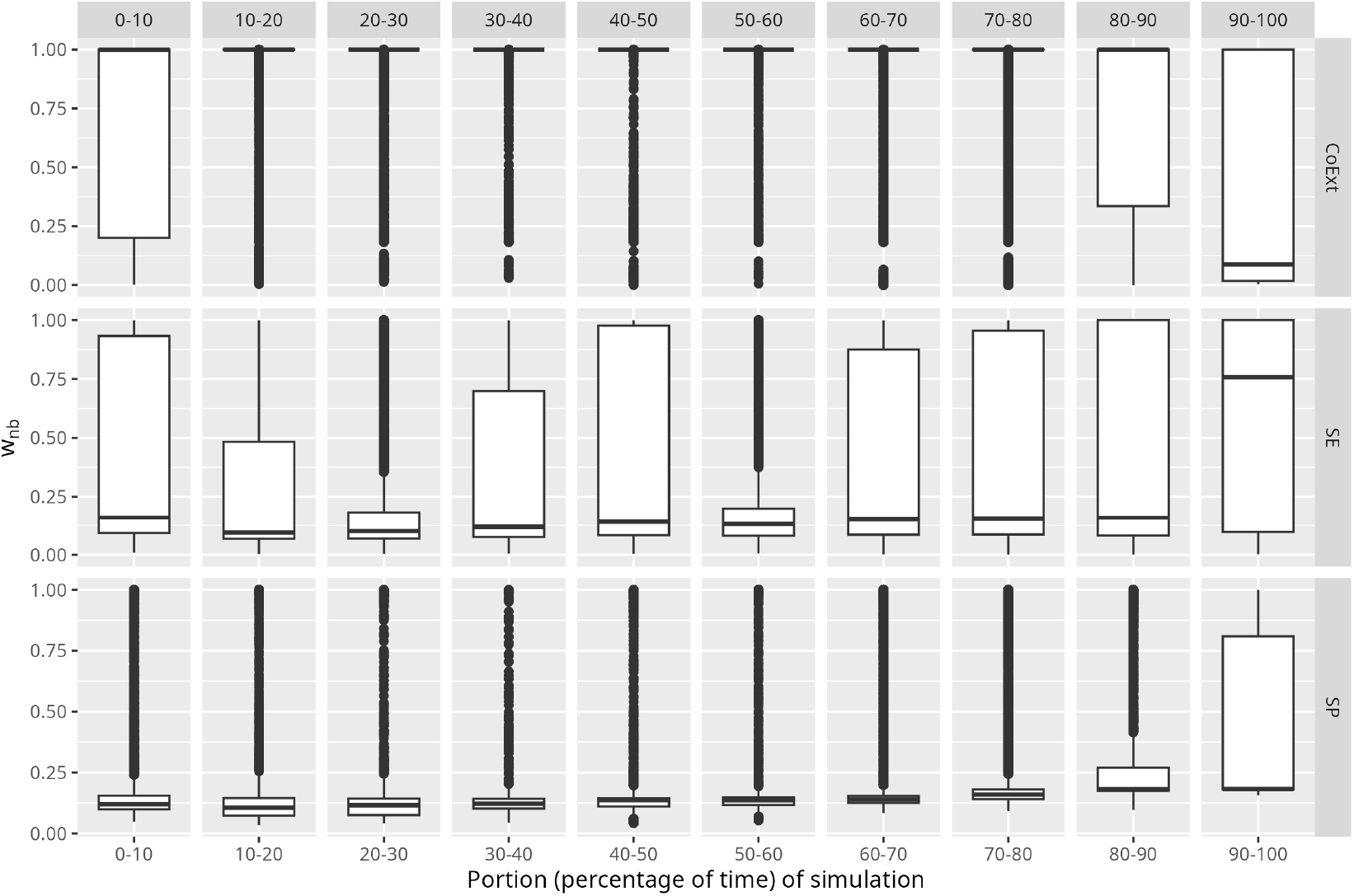
Akaike weights evolution of the negative binomial distribution compared to a Poisson distribution. Simulations are split into ten slices to have a glimpse of the dynamic evolution of Akaike weights of the negative binomial distribution (*w*_nb_) using boxplots.

